# Precise, pan-cancer discovery of gene fusions reveals a signature of selection in primary tumors

**DOI:** 10.1101/178061

**Authors:** Donald Eric Freeman, Gillian Lee Hsieh, Jonathan Michael Howard, Erik Lehnert, Julia Salzman

## Abstract

The extent to which gene fusions function as drivers of cancer remains a critical open question in cancer biology. In principle, transcriptome sequencing provided by The Cancer Genome Atlas (TCGA) enables unbiased discovery of gene fusions and post-analysis that informs the answer to this question. To date, such an analysis has been impossible because of performance limitations in fusion detection algorithms. By engineering a new, more precise, algorithm and statistical approaches to post-analysis of fusions called in TCGA data, we report new recurrent gene fusions, including those that could be druggable; new candidate pan-cancer oncogenes based on their profiles in fusions; and prevalent, previously overlooked, candidate oncogenic gene fusions in ovarian cancer, a disease with minimal treatment advances in recent decades. The novel and reproducible statistical algorithms and, more importantly, the biological conclusions open the door for increased attention to gene fusions as drivers of cancer and for future research into using fusions for targeted therapy.

## Introduction

While genomic instability is a hallmark of human cancers, its functions have only partially been explained. Point mutations and gene dosage effects result from genomic instability, but they alone do not explain the origin of human cancers (Martincorena et al., 2015). Genomic instability also results in structural variation in DNA that creates rearrangements, including local duplications, deletions, inversions or larger scale intra- or inter-chromosomal rearrangements that can be processed into mRNAs that are gene fusions.

Gene fusions are known to drive some cancers and can be highly specific and personalized therapeutic targets; among the most famous are the BCR-ABL1 fusion in chronic myelogenous leukemia (CML), and the EML4-ALK fusion in non-small lung cell carcinoma (Soda et al., 2007; Nowell and Hungerford, 1960). Fusions are among the most clinically relevant events in cancer because of their use to direct targeted therapy and because of early detection strategies using RNA or proteins; moreover, as they are truly specific to cancer, they have promising potential as neo-antigens (Zhang, Mardis and Maher, 2017; Ragonnaud and Holst, 2013; Liu and Mardis, 2017).

Because of this, major efforts by clinicians and large sequencing consortia attempt to identify fusions expressed in tumors. However, these attempts are limited by critical roadblocks: current algorithms suffer from high false positive rates and unknown false negative rates. Thus, heuristic approaches and filters are imposed, including taking the consensus of multiple algorithms or imposing priority on the basis of gene ontologies given to fusion partners. These approaches lead to what third party reviews agree is imprecise fusion discovery and bias against discovering novel oncogenes (Liu et al., 2015; Carrara et al., 2013; Kumar et al., 2016). Both shortcomings in ascertainment of fusions by existing algorithms and using recurrence alone to assess function limit the use of fusions to discover new cancer biology. As one of many examples, a recent study of more than 400 pancreatic cancers found no recurrent gene fusions, raising the question if this is due to high false negative rates or this means that fusions are not drivers in the disease (Bailey et al., 2016). Recurrence of fusions is currently one of the only standards in the field used to assess functionality of fusions, but the most frequently expressed fusions may not be the most carcinogenic (Saramäki et al., 2008); on the other hand, there may still be many undiscovered gene fusions that drive cancer.

Thus, the critical question, “are gene fusions under-appreciated drivers of cancer?”, is still unanswered. In this paper, we provide several contributions that more precisely define and provide important advances to answering this question. First, we provide a new algorithm that has significant improvements in precision for unbiased fusion detection in massive genomics datasets. Our new algorithm, sMACHETE (scalable MACHETE), significantly builds on our recently developed MACHETE algorithm (Hsieh et al., 2017) to discover new gene fusions and pan-cancer signatures of selection. Its algorithmic advance over MACHETE is to use novel modeling to account for challenges brought on by “big data”: statistical modeling to identify false positives and avoid heuristic or human-guided filters that are commonly imposed by other fusion detection algorithms. We have systematically evaluated sMACHETE’s false positive rate, which is much lower than other algorithms, and show that sMACHETE has sensitive detection of gold standard positive controls. Beyond recovery of known fusions, sMACHETE predicts novel fusions, the focus of this paper. These fusions include recurrent fusions, two of which we validate in independent samples, and recurrent 5’ and 3’ partner genes.

The improved precision of sMACHETE has allowed us to address several unresolved questions in cancer biology. First, until now, a large fraction of ovarian cancers have lacked explanatory drivers beyond nearly universal TP53 mutations and defects in homologous recombination pathways. Because TP53 mutations create genome instability, a testable hypothesis is that TP53 mutations permit the development of rare or private driver fusions in ovarian cancers, and the fusions have been missed due to biases in currently available algorithms. We apply sMACHETE to RNA-Seq data from bulk tumors and find that 91% of the ovarian tumors we screened have detectable fusions and that 54% of the ovarian cancer tumors express gene fusions involving kinase pathways or known Catalogue of Somatic Mutations In Cancer (COSMIC) genes (Forbes et al., 2014). We also identify novel although low-prevalence recurrent fusions in other cancers, including pancreatic cancer, where they have not been described previously.

Frequent recurrence of gene fusions is a hallmark of a selective event during tumor initiation, and this recurrence has historically been the only evidence available to support that a fusion drives a cancer. While private or very rare gene fusions are beginning to be considered as potential functional drivers (Latysheva and Babu, 2016), the high false positive rates in published algorithms prevent a statistical analysis of whether private or rare gene fusions reported exhibit a signature of selection across massive tumor transcriptome databases, such as TCGA. Signatures of selective advantage of fusion expression include recurrent use of a 5’ or 3’ partner, or enrichment of gene families such as those in Catalogue Of Somatic Mutations In Cancer (COSMIC). We formulate and provide the first such analysis.

In sum, sMACHETE is an advance in accuracy for fusion detection in massive RNA-Seq data sets. The algorithm is reproducible and publicly available, and its results have important biological implications. sMACHETE, applied to hundreds of TCGA RNA-Seq samples, in conjunction with new statistical analysis reveals a signature of fusion expression consistent with the existence of under-appreciated drivers of cancer, including selection for rare or private gene fusions in human cancers with implications from basic biology to the clinic.

## Results

### sMACHETE is a new statistical algorithm for gene fusion discovery

We engineered a new statistical algorithm, the scalable MACHETE (sMACHETE), to discover and estimate the prevalence of gene fusions in massive numbers of data sets. The major computational infrastructure of sMACHETE includes a fusion-nomination step performed by the MACHETE. However, sMACHETE includes key innovations mainly focused on controlling false positives arising from analysis of massive RNA-Seq data sets for fusion discovery, a problem conceptually analogous to multiple hypothesis testing via p-values but which cannot be solved by direct application of common FDR controlling procedures.

The workflow of sMACHETE is as follows: MACHETE is first run on a subset of samples (the “discovery set”) for fusion discovery and modeling. Models of the effect of sequence composition and gene abundance in generating false positive fusion nomination are applied (Supplemental File). Next, the prevalence of the nominated fusions is efficiently tested in the discovery set along with an arbitrarily large number of added samples (the “test set”), easily numbering thousands, using Sequence Bloom Trees (SBTs; Solomon and Kingsford, 2016) and subsequent statistical modeling (see Fig. 1, Methods and Supplemental File). This step further decreases false positive identification of fusions beyond those decreases achieved by MACHETE, which are already lower than any other published algorithm (Hsieh et al., 2017), and increases the precision of fusion prevalence rate estimation. Intuitively, this step checks whether the prevalence of fusions found by running MACHETE is statistically consistent with the estimated prevalence using a string-query based approach (such as SBT). We note that because the SBT searches for fusion-junctional sequences, samples could be positive for a fusion by a SBT yet negative by MACHETE, which requires spanning reads to nominate fusions (Hsieh et al., 2017).

**Figure 1:**
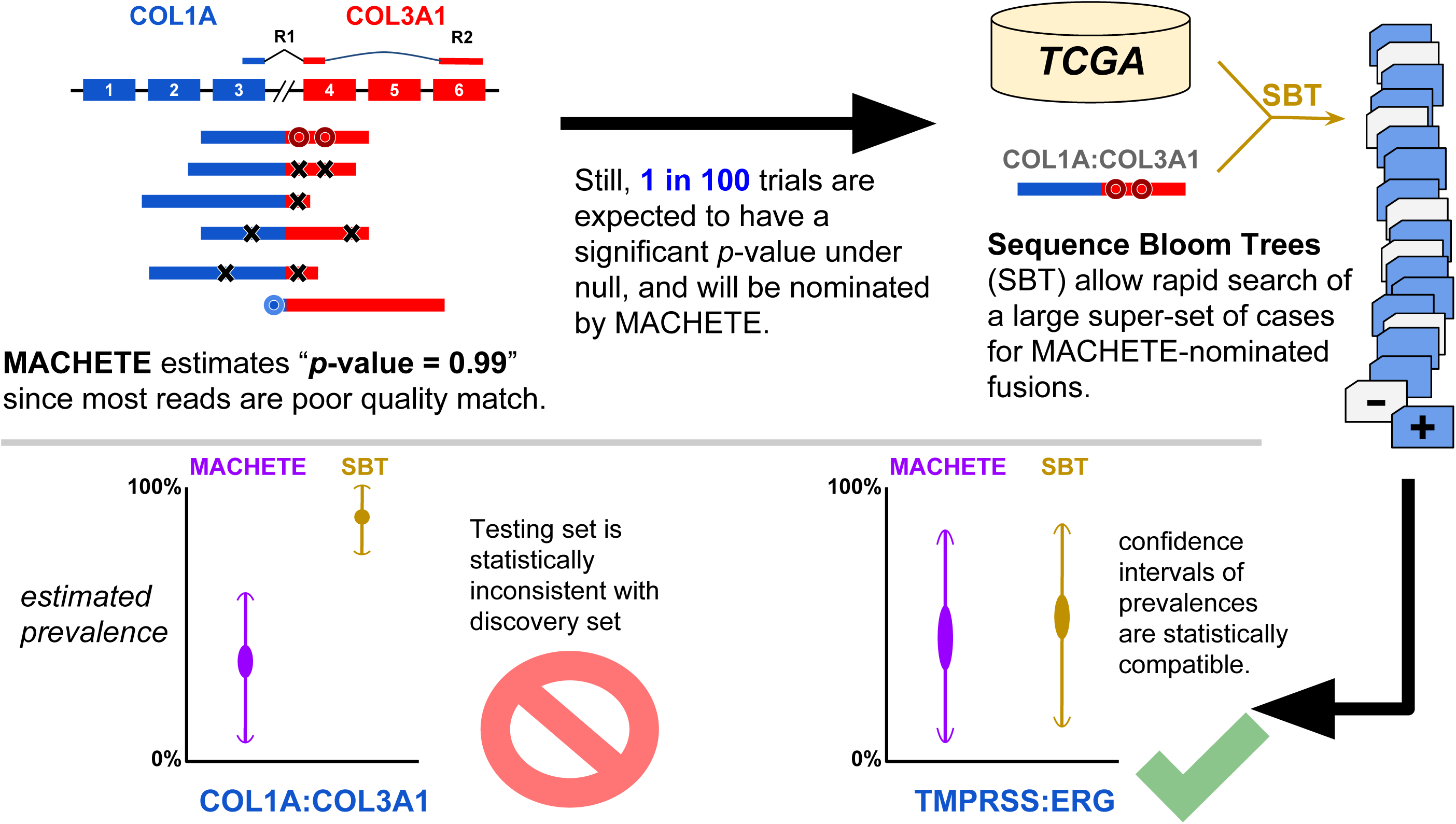
Origin of false positives from MACHETE running on hundreds of data sets. Top left: MACHETE is designed to use all reads, including those censored by other algorithms, to generate an empirical p value for each candidate fusion, computed for each data set separately (Hsieh et al., 2017). Multiple hypothesis testing will result in some fusions passing statistical thresholds under the null. If a single fusion in a single sample has a significant p-value, the sequence will be queried by a SBT which does not use statistical models, and the fusion could be falsely found to be very prevalent. Using confidence intervals based on sampling depth in the discovery and testing sets, analysis of the the SBT can identify false positives (Supplemental File).

**Figure 2:**
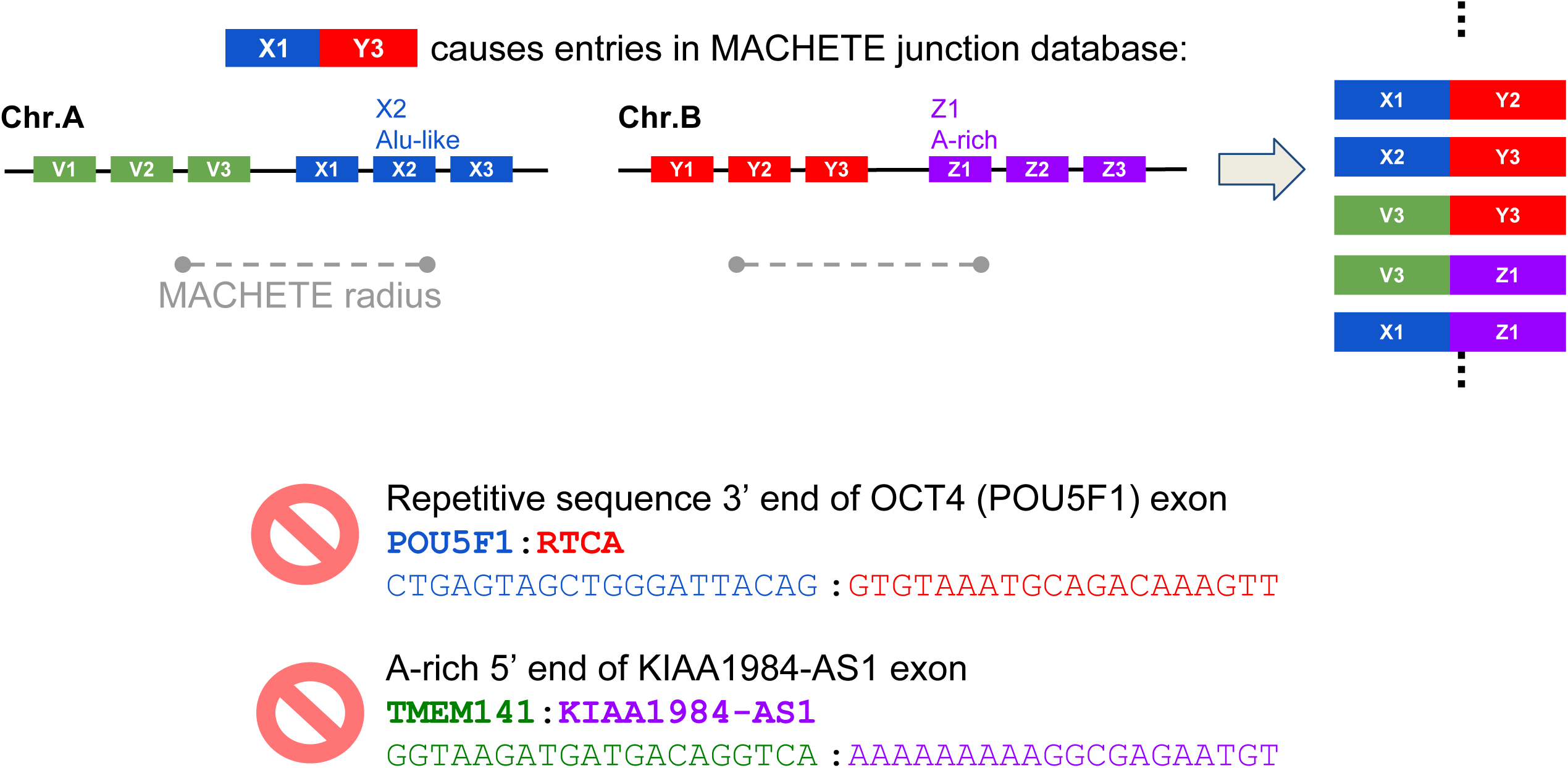
cDNA or mapping artifacts result in inclusion of exon-exon junctions from all permutations of exons within a fixed genomic radius of X1 with all exons in the radius of Y3 in the MACHETE index. Some such exon junctions will include degenerate sequences (left). Because degenerate sequences cannot be mapped uniquely, sMACHETE blinds itself to detection of fusion RNA containing such highly degenerate sequences (for example, due to Alu exonization) or with poly(A) stretches at the 5’ end.

Like MACHETE, sMACHETE does not require human guidance and is a fully automatic pipeline. Moreover, most parts are very portable as they are Dockerized, and most components of the workflows can be easily exported to many platforms using a description given by the Common Workflow Language (CWL). sMACHETE can be applied to any RNA-Seq dataset, including any massive cancer genomics datasets. And, assuming one has access to the secure TCGA database, the analysis we present in this paper is reproducible. (See Supplemental File; also, the code used, including CWL code and Dockerfiles, is available at github sites given in the Supplemental File.)

### sMACHETE improves specificity of fusion detection without sacrificing sensitivity

Compared to current state of the art fusion callers, sMACHETE reduces false positives, the most measurable metric for errors. But this rate can only be exactly computed under simulated conditions where the ground truth is known. As a proxy for ground truth, normal controls are used under the assumption that fusions detected in normal tissues should be rare, as is the case for some germline fusions such as TFG-GPR128 (Chase et al., 2010). We have adopted the common simplifying assumption that prevalent fusions called in normal samples that cannot be explained by readthrough transcription are false positives (Lee et al., 2017; Kumar et al., 2016).

MACHETE, the workhorse of sMACHETE, has been benchmarked on a group of normal samples and simulated data with the lowest false positive rate and highest positive predictive value (PPR) of published algorithms (Hsieh et al., 2017, and Supplemental File). Theoretical analysis of the algorithm formally implies that sMACHETE maintains or improves the already low false-positive rate of MACHETE. In this paper, we go further and quantify sMACHETE’s FPR on the Illumina Body Map data set because it is comparable in its age, depth and read length to TCGA data; further, there are not large numbers of normal samples of the same vintage as the TCGA data analyzed here, and TCGA samples classified as normal are not molecularly normal (personal communication with TCGA). sMACHETE increases specificity on the Body Map compared to the consensus best existing algorithm tested, ChimerSeq (Lee et al., 2017), which reports significantly more fusions in cancer samples that are also detected in normals, suggesting they are false positives (Fig. 4). We have used fusions called by ChimerSeq to compare sMACHETE’s sensitivity and specificity because ChimerSeq entails performance benchmarking of multiple ‘top performing’ algorithms, and, using a disciplined procedure for evaluating them, instantiates a meta-caller to produce more reliable calls than any algorithm independently (Lee et al., 2017).

**Figure 4:**
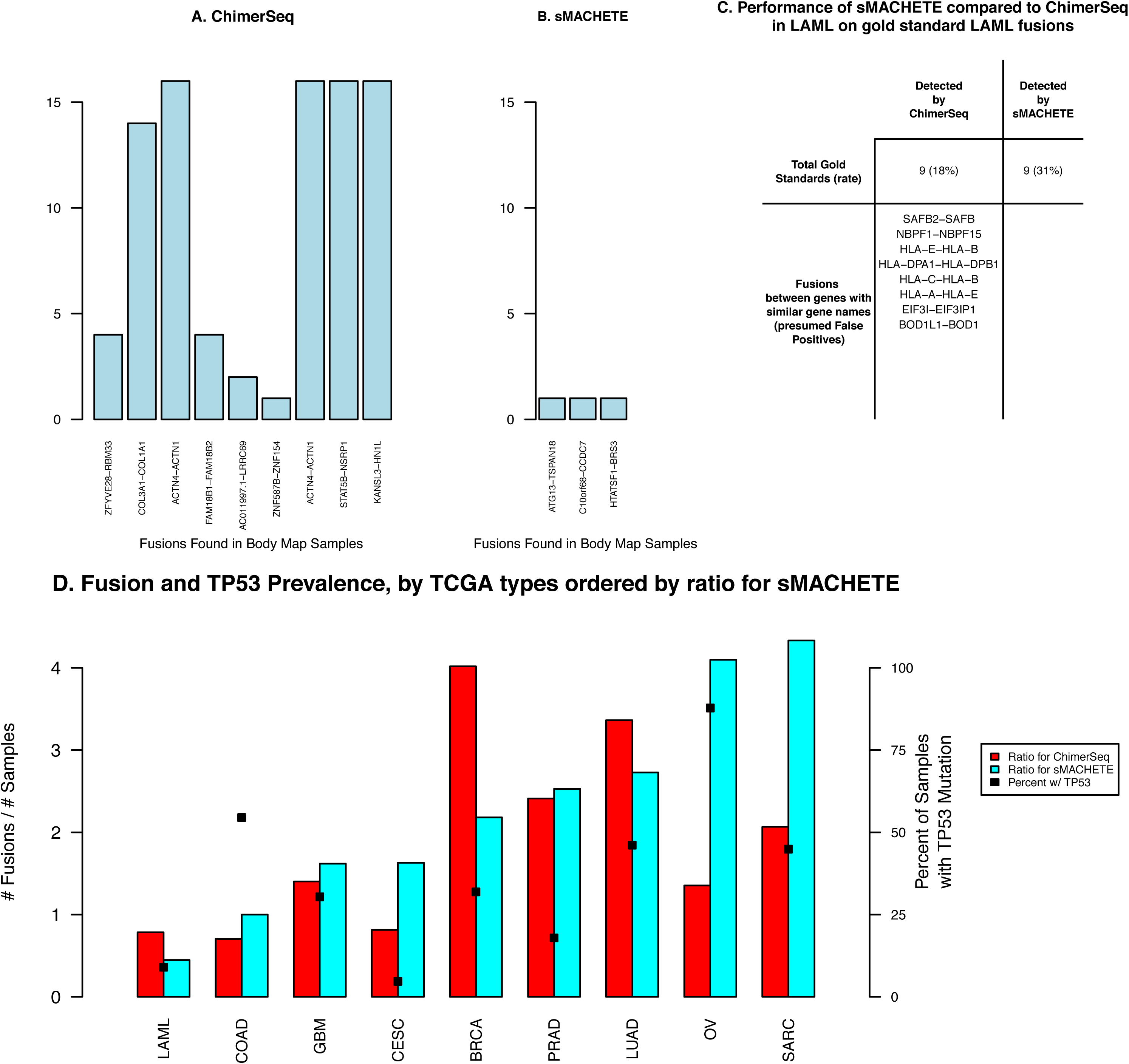
(A) and (B): 9 unique fusions called by ChimerSeq are also detected in Body Map samples by a SBT query, some in all Body Map samples, whereas only 3 fusions are called by sMACHETE in TCGA tumors, and then found in Body Map samples by a SBT query, and only one in each sample. All three of these fusions are intrachromosomal, a feature not true of six of the fusions called by ChimerSeq; (C): Performance of sMACHETE compared to ChimerSeq in LAML: Each algorithm identifies the same number of gold standard LAML fusions, but among likely false positives, ChimerSeq detects 8 while sMACHETE detects none (Supplemental File); (D): Unique fusions identified across all samples in each TCGA disease type per total samples analyzed by sMACHETE. While achieving a significantly lower false positive rate, sMACHETE has improved sensitivity in some diseases with fractions of fusions detected that are more consistent with fraction of TP53 mutations in each disease as reported by cBioPortal (Gao et al., 2013).

Any algorithm’s FPR can be trivially reduced by sacrificing sensitivity. However, we find that sMACHETE’s precision may in fact improve sensitivity. In primary tumors, no ground truth is known, so we use well-studied and generally cytogenetically simple tumor types such as acute myeloid leukemia (LAML) as a best approximation. In a large cohort of LAML samples investigated through both next-generation sequencing and cytogenetics by a large consortium (Cancer Genome Atlas Research Network, 2013; Papaemmanuil et al., 2016), sMACHETE improves the rate of true positive recovery compared to ChimerSeq (Lee et al., 2017), when using nomination of fusions between homologous genes as a proxy for false positives (Fig. 4C, and Supplemental File).

sMACHETE maintains high precision in a variety of solid tumors that have more complex cytogenetics than LAML. This cytogenetic complexity could result in either more false positives or false negatives, as occurs with other algorithms (Stransky et al., 2014; Yoshihara et al., 2015; Van Allen et al., 2016). As one computational test of this, we used the principle of cancer biology that the total number of fusions detected should be correlated with an orthogonal measure of a tumor’s genome stability, as measured by the mutation rate of TP53 (Forment et al., 2012). sMACHETE has much higher correlation with TP53 mutation rate and number of fusions identified per sample compared to the current best performing published fusion caller, ChimerSeq, across tumor types (Pearson correlation .6 and .06 respectively; Spearman rho .45 and .07 respectively; Fig. 4D); and in general calls more fusions in tumors with high TP53 mutation rates, and fewer than ChimerSeq in less cytogenetically complex tumors while retaining tight control of false positives in other samples.

ChimerSeq and sMACHETE report similar numbers of fusions in the same TCGA cohort of the 278 samples that were analyzed in common (Supplemental File). The set of fusions (counted as unique gene pairs, ignoring splice variants) on this set of samples has little overall concordance: 660 unique fusions are called by ChimerSeq, 525 unique gene pairs expressed as fusions are called on this set by sMACHETE, and only 213 are common to both algorithms. Of note, among this list, 8 distinct gene fusions involving HLA or ribosomal protein subunit genes, proxies for likely false positives due to their high expression, are called by ChimerSeq, while none are called by sMACHETE. ChimerSeq appears to call no fusions for, and presumably does not analyze, pancreatic adenocarcinoma (PAAD) tumors. (In our discussion of other tumor types in this paper, we use abbreviations following TCGA nomenclature. See Fig. 3.)

**Figure 3:**
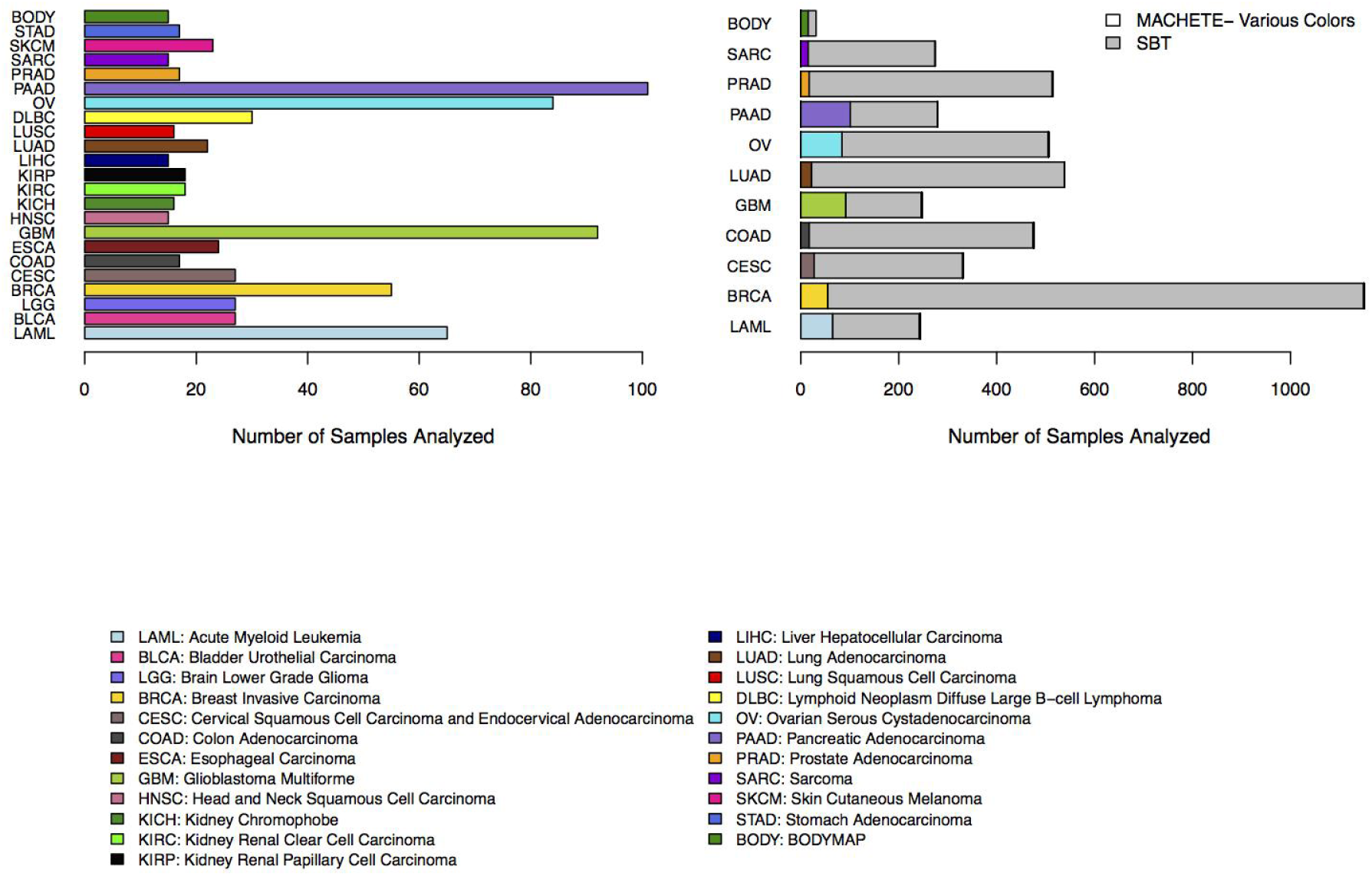
Left panel: Total runs per cancer type in the sMACHETE discovery set. Right panel: Number of cancers in discovery set and in Sequence Bloom Trees for those cancers with Sequence Bloom Trees built.

Because the ground truth is not known for most tumors profiled in the TCGA data, we have investigated the performance on sMACHETE for a handful of well known recurrent gene fusions beyond LAML. As an example, TMPRSS2-ERG is the most commonly known recurrent gene fusion in any solid tumor (Maher et al., 2009). We hand-picked 15 prostate cancer tumors that were positive for TMPRSS2-ERG, as reported in Sadis et al. (2013), to include in the discovery set. sMACHETE detected 7 splice variants of TMPRSS2-ERG, increasing the sensitivity of detecting alternative splice variants of fusions and total prevalence of detected fusions compared to ChimerSeq (Supplemental File and Lee et al., 2017; Gorohovski et al., 2017). The prevalence of TMPRSS2-ERG in prostate adenocarcinoma (PRAD) (Tomlins et al., 2008) detected by sMACHETE and ChimerSeq is similar (39% by sMACHETE and 42% by ChimerSeq).

### sMACHETE predicts novel recurrent fusions validated in independent clinical samples

Apart from sMACHETE’s rediscovery of well-known recurrent gene fusions, the vast majority of sMACHETE’s predicted fusions were present in only a small number of tumors (see Fig. 5 and Supplemental Table 1). While generally low prevalence, several novel fusions were detected at sufficient frequency that they would be expected to appear in an independent, moderate number of primary tumor samples that our laboratory could reasonably test.

**Table 1:**
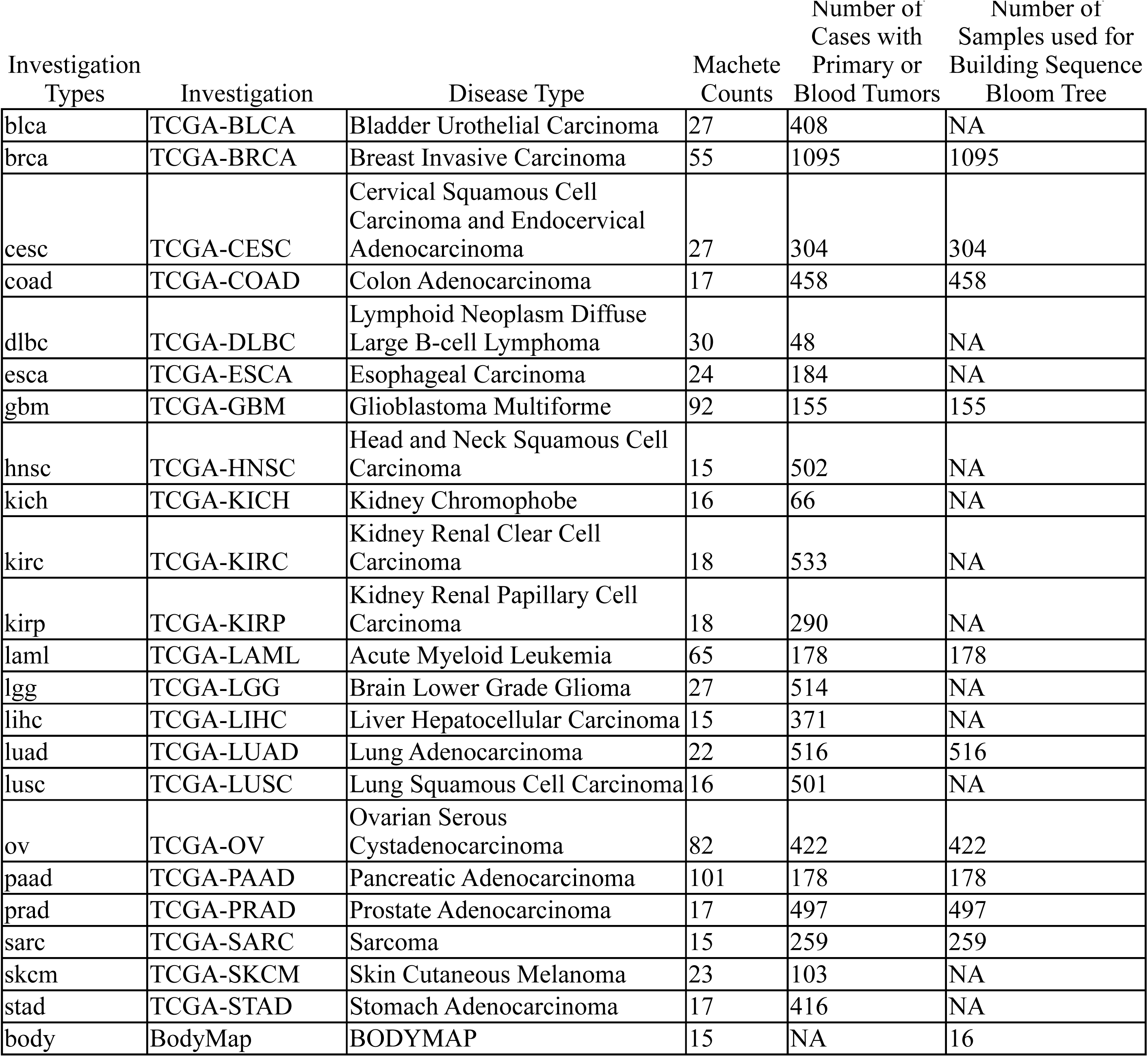
Number of Samples Analyzed by MACHETE and sMACHETE, and Total Number of Cases and Samples in TCGA Data Set

| Investigation Types | Investigation | Disease Type | Machete Counts | Number of Cases with Primary or Blood Tumors | Number of Samples used for Building Sequence Bloom Tree |
| --- | --- | --- | --- | --- | --- |
| blca | TCGA-BLCA | Bladder Urothelial Carcinoma | 27 | 408 | NA |
| brca | TCGA-BRCA | Breast Invasive Carcinoma | 55 | 1095 | 1095 |
| cesc | TCGA-CESC | Cervical Squamous Cell Carcinoma and Endocervical Adenocarcinoma | 27 | 304 | 304 |
| coad | TCGA-COAD | Colon Adenocarcinoma | 17 | 458 | 458 |
| dlbc | TCGA-DLBC | Lymphoid Neoplasm Diffuse Large B-cell Lymphoma | 30 | 48 | NA |
| esca | TCGA-ESCA | Esophageal Carcinoma | 24 | 184 | NA |
| gbm | TCGA-GBM | Glioblastoma Multiforme | 92 | 155 | 155 |
| hnsc | TCGA-HNSC | Head and Neck Squamous Cell Carcinoma | 15 | 502 | NA |
| kich | TCGA-KICH | Kidney Chromophobe | 16 | 66 | NA |
| kirc | TCGA-KIRC | Kidney Renal Clear Cell Carcinoma | 18 | 533 | NA |
| kirp | TCGA-KIRP | Kidney Renal Papillary Cell Carcinoma | 18 | 290 | NA |
| laml | TCGA-LAML | Acute Myeloid Leukemia | 65 | 178 | 178 |
| lgg | TCGA-LGG | Brain Lower Grade Glioma | 27 | 514 | NA |
| lihc | TCGA-LIHC | Liver Hepatocellular Carcinoma | 15 | 371 | NA |
| luad | TCGA-LUAD | Lung Adenocarcinoma | 22 | 516 | 516 |
| lusc | TCGA-LUSC | Lung Squamous Cell Carcinoma | 16 | 501 | NA |
| ov | TCGA-OV | Ovarian Serous Cystadenocarcinoma | 82 | 422 | 422 |
| paad | TCGA-PAAD | Pancreatic Adenocarcinoma | 101 | 178 | 178 |
| prad | TCGA-PRAD | Prostate Adenocarcinoma | 17 | 497 | 497 |
| sarc | TCGA-SARC | Sarcoma | 15 | 259 | 259 |
| skcm | TCGA-SKCM | Skin Cutaneous Melanoma | 23 | 103 | NA |
| stad | TCGA-STAD | Stomach Adenocarcinoma | 17 | 416 | NA |
| body | BodyMap | BODYMAP | 15 | NA | 16 |

**Figure 5:**
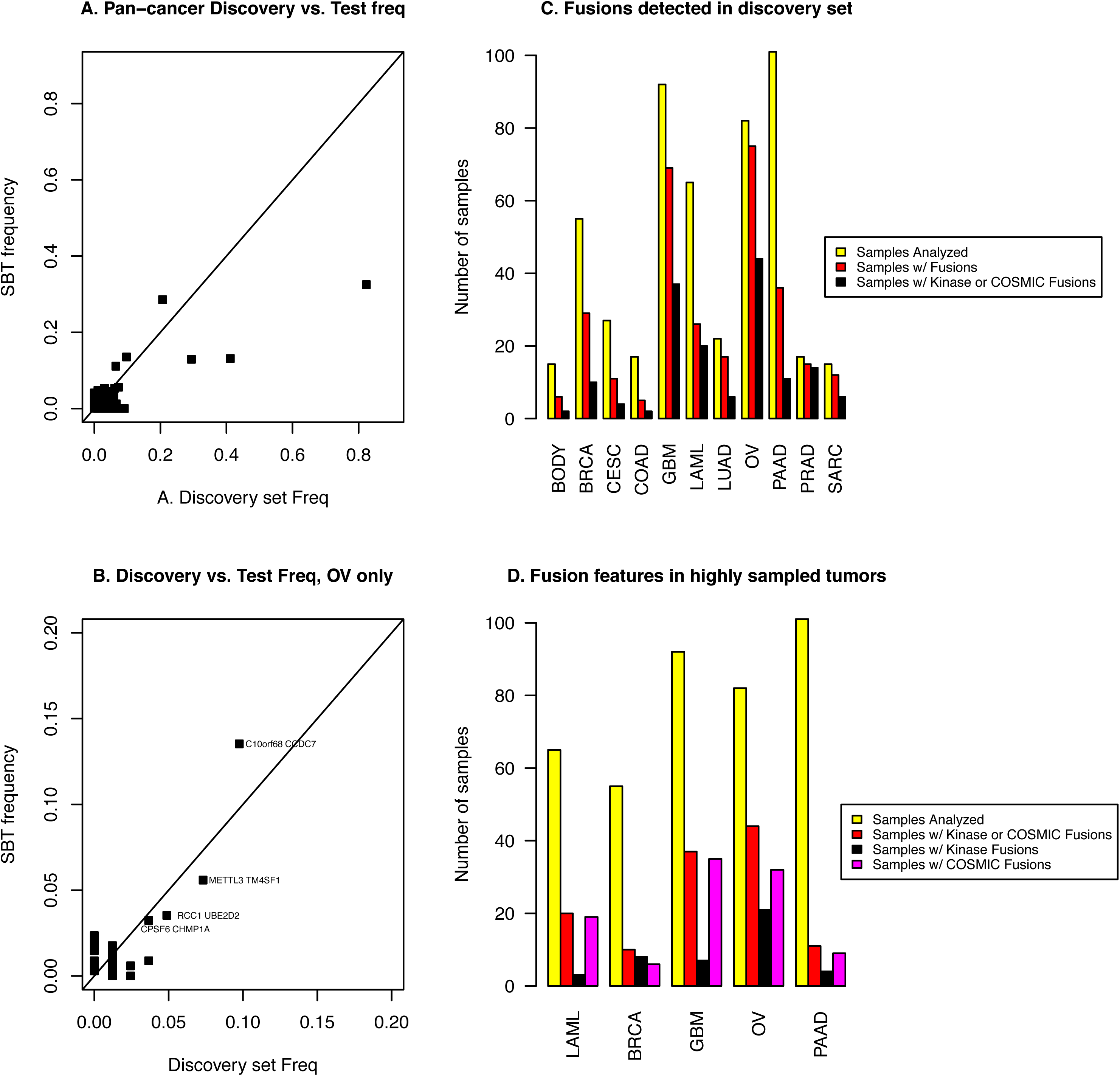
(A) and (B:) Relationship between estimated fusion prevalence between discovery set and test set as quantified by SBT: (A) all fusions and (B): only fusions in ovarian cancer. (C): Rate of fusion detection in discovery set including those fusions annotated to include COSMIC genes and the term kinase; (D) more detailed analysis of highly sampled tumors. ∼90% of ovarian cancers in our discovery set have a sMACHETE-called fusion.

Using sMACHETE’s predictions from TCGA data, we attempted to validate four novel and one previously reported recurrent fusions on nine primary ovarian tumor samples, labeled (A-I). We first tested for two novel fusions: CPSF6-CHMP1A, a fusion consistent with deriving from interchromosomal rearrangement, and RB1-ITM2B, a rearrangement between two neighboring genes. Samples (C,E,F) (33%) had PCR products of the expected size for CPSF6-CHMP1A and samples (B,E,F,G,H,I) (>50%) had PCR products of the expected size for RB1-ITM2B. Sanger sequencing of the PCR products produced the expected sequences (see Figures 6A and 6B, Methods and Supplemental File). RB1-ITM2B could be explained by a cancer-specific circular RNA or a local genomic rearrangement (see Fig. 6A); we have not previously detected this sequence in normal samples (Szabo et al., 2015, Hsieh et al., 2017). While we did not attempt to distinguish whether an underlying DNA change was responsible for the RB1-ITM2B fusion, the estimated prevalence of RB1-ITM2B from poly(A) selected TCGA libraries was only 2%. This is much lower than the 55% prevalence detected by PCR, and is consistent with the hypothesis that RB1-ITM2B is a circRNA that is depleted in poly(A) selected libraries.

**Figure 6:**
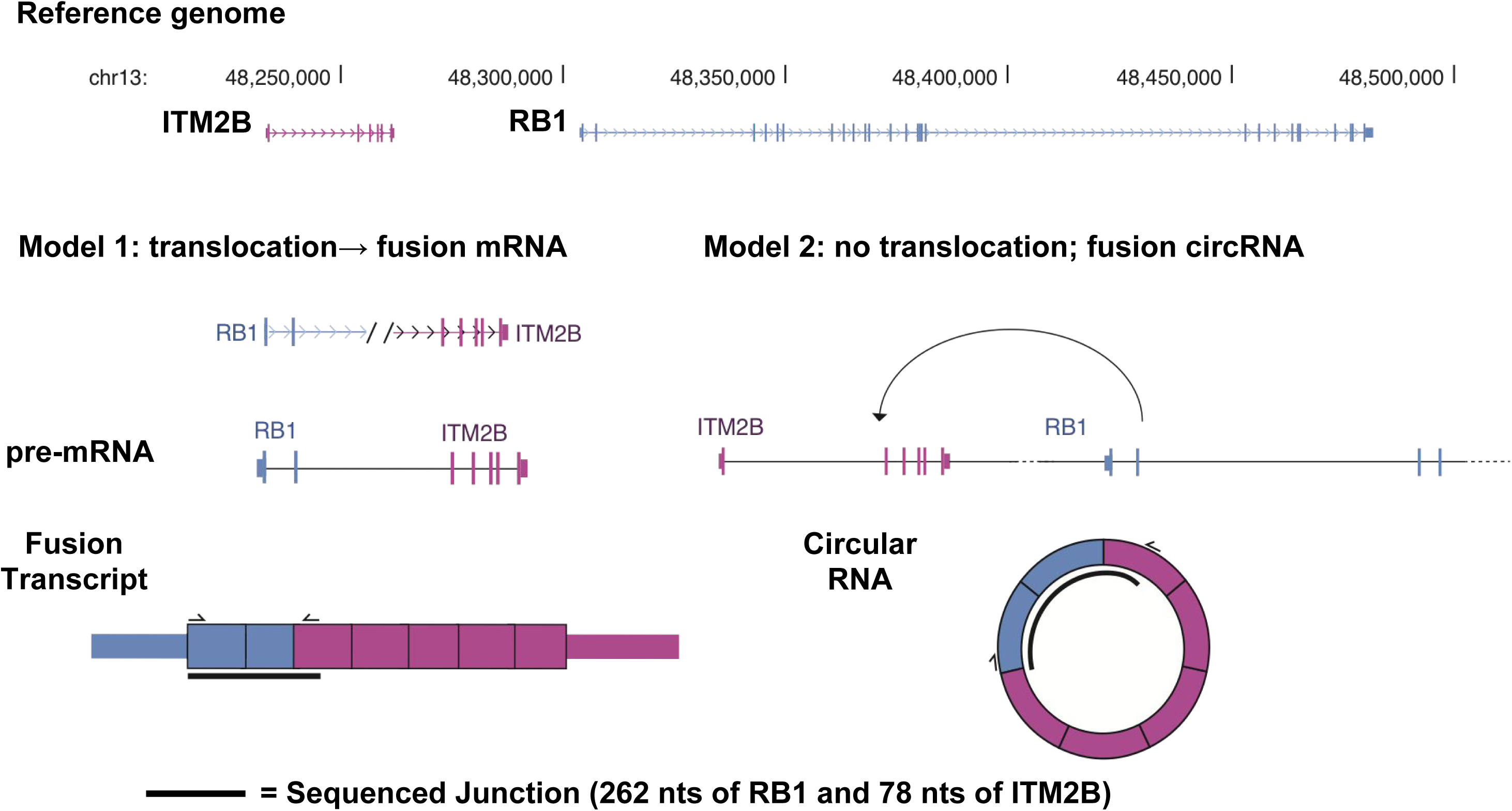
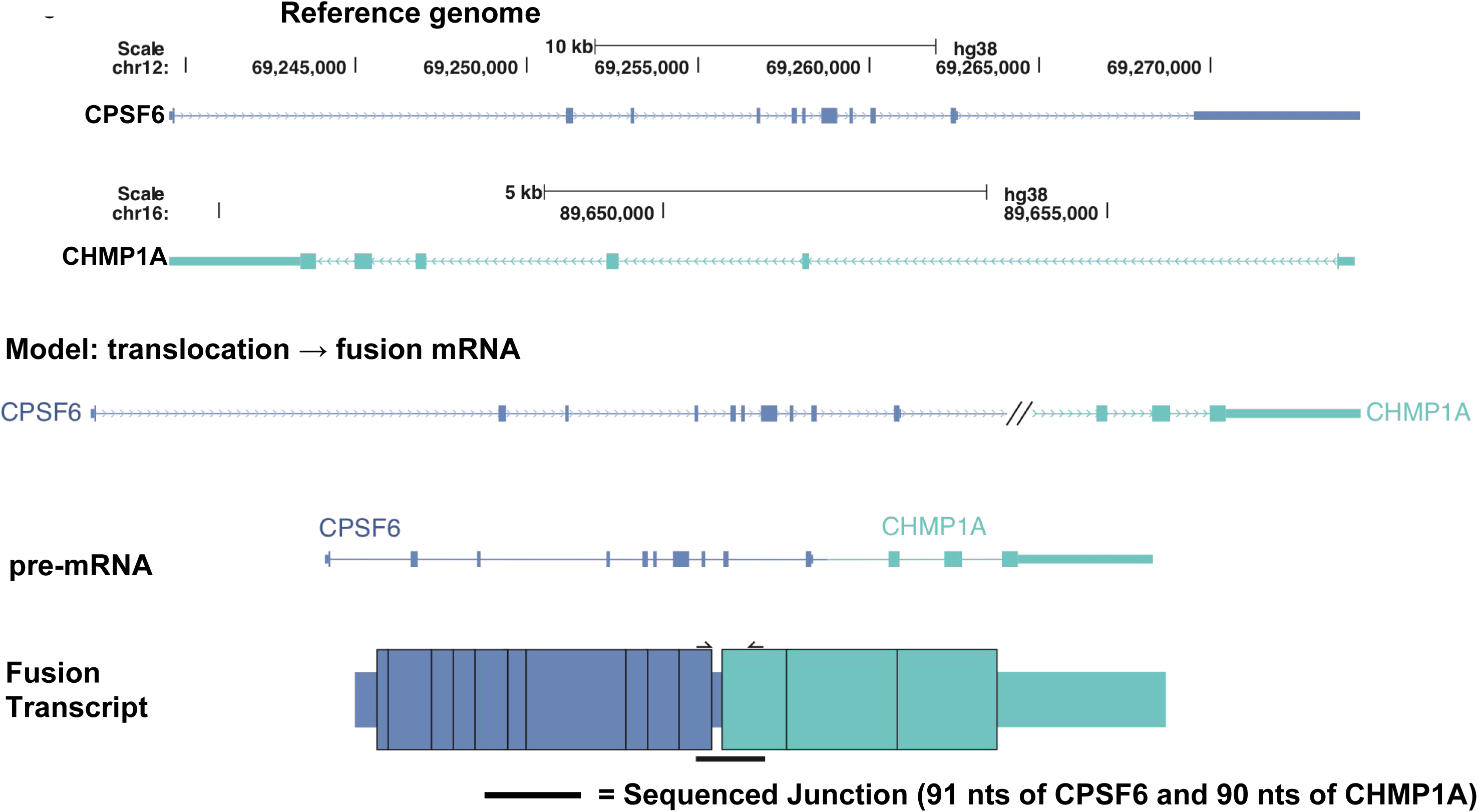
(A) In the reference genome, ITM2B is upstream of RB1 and both genes are transcribed in the sense orientation. In Model 1 (L), a genomic change, such as a tandem duplication, puts the genomic sequence of RB1 upstream of exons of ITM2B. Transcription from the RB1 promoter results in a pre-mRNA that is spliced into a fusion mRNA. In Model 2 (R), no DNA rearrangement occurs, but readthrough transcription from the ITM2B promoter results in a pre-mRNA that is back-spliced into a circRNA containing exons of RB1 and ITM2B. The sequenced junction contains 262 nt of RB1 and 78 nts of ITM2B; (B) CPSF6 is transcribed from chr12 and CHMP1A from chromosome 16. Model for the fusion CPSF6-CHMP1A; sequenced junction contains 91nt upstream of and 90nt downstream of the fusionf junction.

We tested the same samples for three other fusions detected by sMACHETE: a previously known germline fusion, TFG-GPR128 (Chase et al., 2010) and two predicted ovarian-specific recurrent fusions, METTL3-TM4SF1 and RCC1-UBE2D2. Consistent with the range of previous reports of the prevalence of TFG-GPR128 in the population (3/120 as reported in Chase et al., 2010, 95% CI: 0.5%-7.1%), sMACHETE estimates its frequency in TCGA data to be <1% in sarcoma (SARC), 2.2% in PAAD, and 1.4% in ovarian serous cystadenocarcinoma (OV) (see Supplemental Table 1). The predicted frequency of METTL3-TM4SF1 and RCC1-UBE2D2 were similarly low (5.9% and 3.8% of OV cases, respectively). All samples tested by PCR for these three fusions were negative, which is consistent with their estimated prevalence under a simple binomial sampling model. Because of the low prevalence, a much larger sample size, greater than one hundred, would be necessary to provide sufficient statistical power to test if these fusions are recurrent.

### Fusions identified by sMACHETE are enriched in known oncogenes

Because we, and the vast majority of researchers, do not have access to TCGA samples for additional PCR validation, we used orthogonal computational tests of sMACHETE’s novel fusion predictions to support the assertion that most of sMACHETE’s fusion predictions are not artifacts. We first investigated the distribution of functional gene ontologies of reported fusion partners, as these are not used by sMACHETE and so provide an independent test of whether sMACHETE is identifying a potentially important biological signal. To test whether the putative fusions identified by sMACHETE are enriched for genes in known cancer pathways, for each cancer type we tested for enrichment of genes present in the Catalogue of Somatic Mutations in Cancer (COSMIC) database or that include the word “kinase” in their annotation (Forbes et al., 2014; Methods). In six of the ten cancer types profiled by sMACHETE, the fraction of samples with fusions identified and annotated as either COSMIC or kinase exceeds 20%, a rate much greater than expected by chance (Methods and Fig. 5C). Among samples with any fusion reported, the largest enrichment for COSMIC or kinase annotated genes are in PRAD (93%) and LAML (77%), as expected because the most frequent gene fusions in PRAD involve the ETS family of transcription factors (COSMIC genes), and LAML is a disease where fusions have been intensively studied, include known drivers, and whose partners are therefore annotated as COSMIC genes (Forbes et al., 2014; Fig. 5).

### Ovarian cancers have high fusion prevalence and are enriched kinase and COSMIC genes

The most common genetic lesion in ovarian cancer is the TP53 mutation, present in 88% of cases (cBioPortal, retrieved July 18, 2017, see Gao et al., 2013), although there is debate in the literature that this prevalence is an underestimate. Regardless, other drivers must exist because, for example, TP53 mutations are not sufficient to cause cancers (Martincorena et al., 2015). In OV, such explanatory driving events are as yet unknown (Bowtell et al., 2015). The prevalence of TP53 mutations generates the hypothesis that the resulting genome instability could generate fusions responsible for driving some fraction of these cancers, but which have been missed because of shortcomings in other available algorithms; we sought to test this hypothesis.

sMACHETE reports 91% of all ovarian cancers in its discovery set to have a gene fusion, the highest rate of any disease we profiled. 54% of ovarian tumors contain a fusion involving a kinase or COSMIC gene, a higher frequency than any other profiled disease (see Fig. 5). Prevalent recurrent fusions were not detected, with the exception of one that is most parsimoniously explained by being circRNA: a putative fusion between C10orf68 and CCDC7, a pair of genes with overlapping transcriptional boundaries and shared exons, one of only two fusions called in both our Body Map and tumor samples (Supplemental Table 1). This fusion is also reported in LAML by a separate group, and is consistent with the fusion being a circular RNA (Cancer Genome Atlas Research Network, 2013).

Recurrent fusions of low prevalence involving genes on different chromosomes, unlikely to be circRNA, were detected as described above: 3.8% of tumors were estimated to have the fusion RCC1-UBE2D2. RCC1 is a regulator of chromosome condensation and UBE2D2 is an ubiquitin conjugating enzyme. RCC1-UBE2D2 is predicted to be specific to ovarian tumors. The fusion METTL3-TM4SF1 of METTL3, a methyltransferase-like protein involved in splicing, and TM4SF1, a transmembrane protein of unknown function, was seen in 5.9% of tumors and also specific to ovarian cancer.

sMACHETE predicts that the rate that fusions are present in ovarian cancer is higher than previously reported by other analyses of TCGA data (Yoshihara et al., 2015; Earp et al., 2017). To be called by sMACHETE, a fusion must be nominated by MACHETE. Thus, the comprehensive tests of MACHETE’s false positive rates in Hsieh et al. (2017) imply a low false positive rate for sMACHETE. This, together with the results in this paper, argue against the possibility that sMACHETE’s discoveries are due to ‘lax controls‘ on false positive rates and instead strongly suggest a biological differentiation of ovarian cancer fusion expression from other cancers we profiled. The enrichment of COSMIC genes in fusion partners further supports this.

Further, our discovery of a high fraction of gene fusions in ovarian cancer is consistent with an orthogonal metric of genome instability in this disease, its TP53 mutation rate of 88% (Methods; TCGA, 2011). This, along with sMACHETE’s specificity on normal controls, supports the interpretation that fusions, perhaps relatively rare or private events, could be an unappreciated driver of ovarian cancers (see Fig. 5). Functional tests of this hypothesis are important but beyond the scope of this paper, and there is an important clinical implication that if rare or low prevalence fusions are common, and if some are potentially druggable, then ‘personalized’ tumor profiling would be needed to inform treatment.

### Statistical analysis of private fusions predicts new oncogenes

Fusions that recur with relatively high frequencies across cases are appreciated to have a selective advantage for tumors, because recurrence has historically been used as a proxy for function in cancer biology. However, statistical signals in rare fusions, including private fusions that are observed only once, could still have statistical features that distinguish them from molecular events deemed ‘passengers’. While intuition for this idea has been appreciated (Lin et al., 2016; Latysheva et al., 2016), statistical formalism has been missing. Mathematical modeling shows that such private fusion expression is, somewhat counter-intuitively, to be expected in the 739 cases we profiled if a moderate fraction of human genes could function as oncogenes when participating in fusions (Supplemental File). Intuitively, this is because of quadratic growth in the number of possible combinations of fusions if a group of genes can serve as oncogenic 5’ or 3’ partners, which implies very high sampling may be required to observe recurrence.

A large number (660) of the 1006 gene fusions (760 unique gene fusions, as some occur multiple times) identified by sMACHETE in the TCGA tumor set are observed only once in our set of profiled tumors (i.e., they are private). (The number 660 is a numerical coincidence with the 660 reported earlier regarding fusions called by ChimerSeq.) We tested whether the 5’ or 3’ partners reappeared on the list of private fusions more often than would be expected compared to a null distribution using a statistical model that is a generalization of the well-known “birthday problem” (Henze, 1998, Supplemental File). We omitted recurrent fusions in the analysis of enrichment for 5’ and 3’ partners as a conservative measure to prevent a bias for re-discovering known oncogenic fusions and enriching a statistical signal, because many gene fusions that are recurrent have had functional assignments as oncogenes because there is bias towards studying them.

This analysis establishes both the excess or ‘effect size’ for the number of genes recurrently present in a 5’ and 3’ fusion and statistical significance (Supplemental File). sMACHETE reports 38 recurrent 5’ partners and 33 recurrent 3’ partners, with both having corresponding p-values << 10^-5^, which are highly statistically significant findings. Moreover, this is a finding with a large effect size: sMACHETE predicts tens of novel oncogenic fusion partners from this analysis, which is based on profiling completely private gene fusions; deeper sequencing or larger sample sizes and more cases or cancer types could further increase this number.

In principle, any gene fusion, including recurrent gene fusions, may be expressed due to a predisposition for genomic rearrangement between two loci rather than RNA expression conferring a particular advantage to the tumor. Thus, in addition to the above statistical evidence, we investigated the gene ontology of genes with multiple partners using the logic that gene fusions can activate oncogenes through a variety of mechanisms, for example those that result in omission of a functional domain through truncation (Shirole et al., 2016) that could have similar effects to point mutations. If our analysis is identifying a real signal, we expect some known oncogenes should be reidentified and enriched as gene partners identified in the above analysis.

We find that known oncogenes are amongst the most significantly enriched 5’ and 3’ partners in private gene fusions. For example, RALA, a Ras-family G-protein and known oncogene (Lim et al., 2005), has three distinct partners found in OV and GBM; to our knowledge it has not been previously reported as a recurrent fusion partner, a feature suggesting that it functions as an oncogene through gene fusion. A fourth fusion involving RALA, RALA-YAE1D1, was identified by sMACHETE as a recurrent gene fusion in OV (see Supplemental Table 1), and hence did not contribute to RALA’s score by this method. ZBTB20, a known oncogene (Lim et al., 2005; Zhao, Ren, and Tang, 2014), was also recovered purely on the basis of participating in private fusions. SORL1 (Uren et al., 2008), a putative oncogene, had the highest diversity of 5’ and 3’ partners. UVRAG, a tumor suppressor with activating oncogene activity (He and Liang, 2015), was also found to have multiple partners and has previously not been reported as participating in fusions. Many other genes on sMACHETE’s list had statistical signal consistent with being novel oncogenes (see Supplemental Table 1).

### Pan-cancer analysis reveals novel rare recurrent fusions expressed in multiple cancer types

Classically, recurrent gene fusions have been considered to be specific to particular tumor-types, such as BCR-ABL1 fusions in CML, EWSR1-FLI1 fusions in Ewing’s sarcoma, and TMPRSS2-ERG fusions in prostate cancers. Next-generation sequencing has revealed exceptions to these initial findings, such as the existence of BCR-ABL1 fusions in LAML and the surprising discovery of EWSR1-FLI1 fusions in pancreatic neuroendocrine tumors (Scarpa et al., 2017).

These examples raise the possibility that within a single cancer type (in the above example, LAML or pancreatic neuroendocrine tumors) low-prevalence recurrent gene fusions could be drivers of these specific tumor cases above, and more generally that recurrent fusions that are rare within a tumor type could drive some cancers. In this scenario, either very high sample sizes or pan-cancer analysis would be necessary to detect them. Further, if some of these fusions were recurrent across a pan-cancer panel, but had low overall prevalence, surveys of the TCGA datasets by consortia studying a single tumor may have missed them because such analysis typically involves profiling only one disease. We sought to test if, like private fusions, sMACHETE identified rare recurrent fusions that were observed at rate higher than expected by chance and that would be consistent with being under selection (Supplemental File).

sMACHETE predicted 100 recurrent gene fusions, indeed far more than would be expected by chance (Supplemental File). This list includes fusions detected in more than one cancer and those that involve partners with annotations indicating potential druggability, such as kinases, chromatin remodeling complexes, and other signaling molecules (e.g., Strawberry Notched Homolog, SBNO2, in the putative fusion product SBNO2-SERINC2; Supplemental Table 1). Another example is a fusion involving the ribosomal protein kinase RPS6KB1-VMP1, previously identified as a recurrent fusion in breast invasive carcinoma (BRCA) (Inaki, et al., 2011), which was detected for the first time in other cancer types, such as lung adenocarcinoma (LUAD) and OV (Supplemental Table 1). PAAD, which had previously lacked reports of recurrent fusions, was found to harbor a group of low-prevalence recurrent fusions when all cancer types were used to estimate recurrence. Some of these rare recurrent gene fusions were present across tumor types in addition to PAAD; for example, ERBB2-PPP1R1B was detected in two total tumors across TCGA including once in PAAD. The examples above represent fusions that in principle, could conceivably be targetable with current drugs (Supplemental Table 1), pending further tests. They show the potential for fusions, and not just point mutations, to stratify patients clinically.

## Discussion

Some of the first oncogenes were discovered with statistical modeling that linked inherited mutations and cancer risk (e.g. Knudson, 1971). The advent of high-throughput sequencing has promised the discovery of novel oncogenes which can inform basic biology and provide therapeutic targets or biomarkers (Cibulskis et al., 2013; Lawrence et al., 2014).

However, unbiased, sequencing-based, methodologies for discovery of novel oncogenic gene fusions have been only partially successful. Many likely driving, and druggable, gene fusions have been identified by high-throughput sequencing, but studies reporting them have a non-tested or non-trivial false positive rate even using heuristic or ontological filters, making them unreliable for clinical use. These problems also limit their sensitivity in unbiased screens of massive data sets to discover fusions, novel oncogenes or signatures of evolutionary advantage for rare or private gene fusions.

In this paper, we present sMACHETE, a unified, reproducible statistical algorithm to detect gene fusions in RNA-Seq data set without human-guided filtering. sMACHETE has significantly lower false positive rates than other algorithms. These filters have not sacrificed detection of known true positives. Further, sMACHETE assigns a statistical score that can be used to prioritize fusions on the basis of statistical support, rather than the absolute read counts supporting the fusion. Because of this, like any statistical test, by adjusting the threshold on scoring, sMACHETE’s discovery rate can be tuned to adjust the trade-off between sensitivity and specificity, a feature unavailable in other algorithms but of potential scientific and clinical utility (Hsieh et al., 2017).

The sMACHETE algorithm improves detection of gene fusions that have been missed by other algorithms’ list of “high confidence” gene fusions. Analysis of these gene fusions uncovers new cancer biology: evidence that gene fusions are more prevalent than previously thought in high grade serous ovarian cancers, which lack explanatory oncogenic events, and perhaps are a contributing driver of these cancers. Unlike other algorithms, sMACHETE finds an enrichment of fusions in ovarian cancers that is consistent with the extremely high representation of TP53 mutations in these tumors.

Also, sMACHETE allows for the first rigorous and unbiased quantification of gene fusions in solid tumors, and for tests of whether partners in gene fusions are present at greater frequencies than due to chance. We find positive results, suggesting that gene fusions, even if not recurrent themselves, are under selection by the tumor. Many fusion partners are detected in more than one cancer type, which suggests that fusions may be lesions like point mutations, present across tumors rather than tumor-defining, and suggests that by focusing on one tumor type to detect recurrence, some important cancer biology is lost. Finally, it is also possible that some fusions identified by sMACHETE, especially those that are local, could be germline fusions, passengers or perhaps markers of genetic predisposition for cancer risk, topics we intend to explore further in other work.

While sMACHETE has increased the accuracy of fusion detection, there are two obvious extensions of this work. First, we could include all samples with known, clinically validated fusions in sMACHETE’s discovery set, enabling a strictly higher chance of discovering clinically actionable events. This might further extend the list of potentially druggable fusions that sMACHETE finds. Above, we described fusions between genes where one gene can be drugged by existing therapies, including ERBB2 (HER2/neu). Further work with a clinical focus is needed to determine the extent of potentially druggable fusions identified by sMACHETE, including determinations of whether protein domains targeted by these drugs are included in the fusion. Second, we have limited our analysis to fusion RNAs that occur at annotated exon-exon boundaries; we believe that extending the statistical approaches used to discover gene fusions may allow us to relax the requirement that gene fusions be detected at annotated exonic sequences, without sacrificing the false positive rate. Doing so will provide a more powerful test of whether genomic instability in cancers results in gene fusions that are a “passenger” of this instability or that have currently under-appreciated functional and perhaps clinical importance.

## Methods

### An enhanced statistical framework for large scale genomics

We ran MACHETE on a discovery set of 739 samples from 22 cancers in the TCGA, consisting of a large fraction of LAML (n=65, 37% of individuals represented in the TCGA database), serous ovarian cancer (n=82, 19% of individuals with primary tumors in the database), pancreatic cancer (n=101, 57% of individuals with primary tumors) and glioblastoma (n=92, 59% of individuals with primary tumors) and a small fraction of the other cancers (399 in 18 cancers, 6% of individuals with primary tumors profiled by the TCGA). The remaining samples were designated and used as “testing” data (see Table 1, Supplemental Table 2, Fig. 3 and Supplemental File). As negative controls, we analyzed Illumina Human Body Map data sets (Table 2) because, as described by the TCGA consortium, samples classified as “Solid Tissue Normal” in the TCGA data sets are not consistently molecularly normal. In the discovery step, due to cost limitations, we deeply sampled a subset of tumors; OV, GBM, and PAAD were selected as diseases where early detection or new drug targets could have great impact, and LAML was selected due to its extensively studied cytogenetics.

**Table 2:**
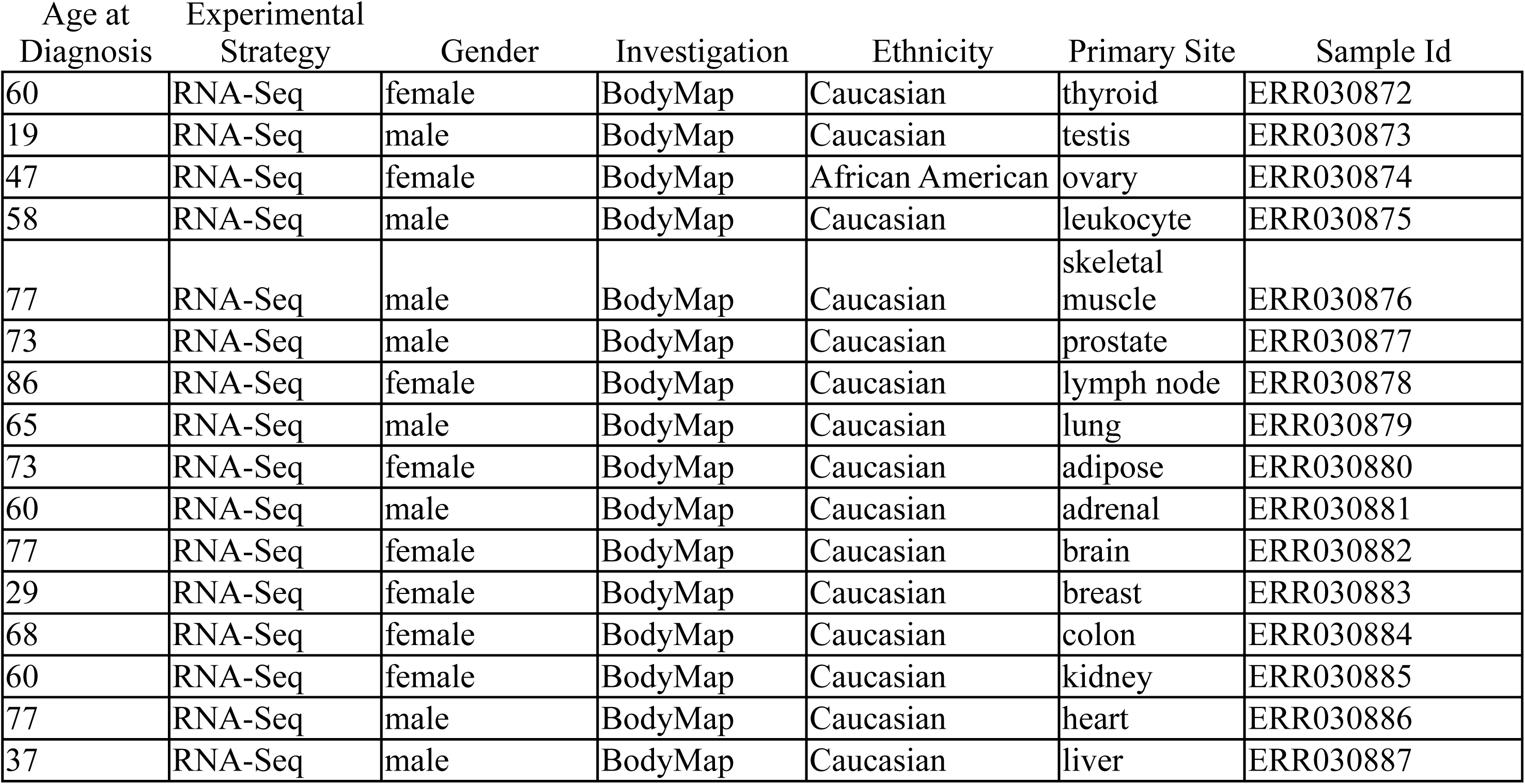
List of All Body Map Samples Used

| Age at<br>Diagnosis | Experimental<br>Strategy | Gender | Investigation | Ethnicity | Primary Site | Sample Id |
| --- | --- | --- | --- | --- | --- | --- |
| 60 | RNA-Seq | female | BodyMap | Caucasian | thyroid | ERR030872 |
| 19 | RNA-Seq | male | BodyMap | Caucasian | testis | ERR030873 |
| 47 | RNA-Seq | female | BodyMap | African American | ovary | ERR030874 |
| 58 | RNA-Seq | male | BodyMap | Caucasian | leukocyte | ERR030875 |
| 77 | RNA-Seq | male | BodyMap | Caucasian | skeletal<br>muscle | ERR030876 |
| 73 | RNA-Seq | male | BodyMap | Caucasian | prostate | ERR030877 |
| 86 | RNA-Seq | female | BodyMap | Caucasian | lymph node | ERR030878 |
| 65 | RNA-Seq | male | BodyMap | Caucasian | lung | ERR030879 |
| 73 | RNA-Seq | female | BodyMap | Caucasian | adipose | ERR030880 |
| 60 | RNA-Seq | male | BodyMap | Caucasian | adrenal | ERR030881 |
| 77 | RNA-Seq | female | BodyMap | Caucasian | brain | ERR030882 |
| 29 | RNA-Seq | female | BodyMap | Caucasian | breast | ERR030883 |
| 68 | RNA-Seq | female | BodyMap | Caucasian | colon | ERR030884 |
| 60 | RNA-Seq | female | BodyMap | Caucasian | kidney | ERR030885 |
| 77 | RNA-Seq | male | BodyMap | Caucasian | heart | ERR030886 |
| 37 | RNA-Seq | male | BodyMap | Caucasian | liver | ERR030887 |

We constructed Sequence Bloom Trees (SBTs) for the Illumina Body Map data and for the RNA-Seq data from each primary tumor from ten cancers with the TCGA dataset: LAML, BRCA, cervical squamous cell carcinoma and endocervical adenocarcinoma (CESC), colon adenocarcinoma (COAD), GBM, LUAD, OV, PAAD, PRAD, and SARC. We queried the SBT with all fusions nominated in the discovery step that passed a statistical threshold (Supplemental File).

We used the discovery set to generate a list of fusions passing MACHETE’s statistical bar (see Supplemental Table 3, Fig. 1), including those fusions nominated by running MACHETE on negative controls from the Body Map. We then queried all data sets for any fusions found in any discovery set (see Fig. 1). We estimated the incidence of each fusion in each sample type (each TCGA disease or Body Map) with SBTs. Next, we used standard binomial confidence intervals to test for consistency of the rate that fusions were present in the samples used in MACHETE’s discovery step and the rate that they were found in the SBT. Fusion sequences that were more prevalent across the entire data set than is statistically compatible with the predicted prevalence from the discovery set were excluded from the final list of fusions (see Fig. 1).

For intuition on why this step is important, consider the scheme in Figure 1: given an exon-exon junction query sequence that could be generated by sequencing errors convolved with gene homology or ligation artifacts, SBTs will not consider the alignment profile of all reads aligning to this junction as MACHETE does, e.g., reads with errors or evidence of other artifacts, because reads with mismatches with the query sequence are by definition censored by the SBT. As a result, the SBT, like other algorithms, can have a high false positive rate due to: (a) false positives intrinsic to the Bloom filters used in the SBT (Solomon and Kingsford, 2016); (b) false positive identification of putative fusions due to events such as depicted in Figure 1, even in the presence of a null false positive rate by the SBT itself (Szabo et al., 2015; Hsieh et al., 2017). False positives as in (b) can arise as follows: if a single artifact (e.g. a ligation artifact between two highly expressed genes) in a single sample passes MACHETE’s statistical threshold in the discovery step, this artifact will be included as a query sequence, and the SBT could detect it a high frequency because the statistical models employed by MACHETE are not used by the SBT (see Fig. 1). Testing for the consistency of the rate of each sequence being detected in the discovery set with its prevalence as estimated by SBTs controls for the multiple testing bias described above (see below and Fig. 1).

See the Supplemental File for more detail about the statistical framework.

### Data availability statement

Access to the data used in this paper is controlled by the NCI and can be requested by following the instructions located at https://gdc.cancer.gov/access-data/obtaining-access-controlled-data.

### MACHETE methodology and Cloud Computing Implementation

The MACHETE algorithm was run on 739 samples from the TCGA database using the Seven Bridges Cancer Genomics Cloud (CGC) platform. For details, see the Supplemental File.

### sMACHETE Methodology: Post-processing of MACHETE output and generation of SBT queries

Technical details of the algorithm and analysis are described in the Supplemental File, and the Supplemental File lists the github sites where the code is available.

### Calculations for fusion, COSMIC and kinase fusion prevalence

For reporting of COSMIC and kinase fusion prevalence in tumors profiled by sMACHETE, SBT reports, for each query sequence passing sMACHETE thresholds, were generated on a per-tumor basis, with a matrix of sample by fusion presence/absence statistics. Samples were included if they were present in the SBT and in the MACHETE discovery set. COSMIC genes and genes annotated as “involved in a kinase pathway” were defined by the annotations in the cancer_gene_consensus.csv file downloaded from the COSMIC website and hg19 RefFlat respectively, implying the chance that a randomly chosen gene would be be annotated with the word ‘kinase’ or found in the COSMIC file is <3%. A gene was defined as having the term “kinase” if its refFlat description included the word “kinase”: 4590 out of 207194 distinct transcript names with products annotated with the word kinase were identified in this refFlat file; there are 595 COSMIC genes, out of all human genes.

### Calculations for expected number of recurrent 5’ and 3’ partners

As a test of the likelihood of observing our results, we employ a statistical model of the probability of observing at most the number of repeated genes that we do observe, under the assumption that the genes in each fusion pair are randomly chosen. For the technical statistical framework, see the Supplemental File.

### File downloads

The following files were downloaded on 12/5/2016 from http://cancer.sanger.ac.uk/cosmic/download using sftp to download it from: /files/grch38/cosmic/v77/cancer_gene_census.csv

Hg19 gene annotations were downloaded from the UCSC genome browser using the refFlat annotation and link: https://genome.ucsc.edu/cgibin/hgTables?hgsid=502825941_NQQWFDm7G51vKlIgkPhbm9a4N3N4&hgta_doSchemaDb=hg19&hgta_doSchemaTable=refLink

The list of COSMIC fusions is at http://cancer.sanger.ac.uk/cosmic/fusion.

Files from ChimeraDB (Lee et al., 2016) were downloaded from http://203.255.191.229:8080/chimerdbv31/mdownload.cdb on 12/11/2016.

Mutation rates of the TP53 locus found in each available cancer subtype (51 subtypes) of the Cancer Genome Atlas database were accessed through the cBioPortal cancer genomics portal (http://www.cbioportal.org), accessed on July 18, 2017. A subset of this data was used to generate Figure 4C, a comparison of TP53 mutation rates to fusion/sample for each cancer subtype.

### Sequence Bloom Tree Methodology

Sequence Bloom Trees (SBTs, Solomon and Kingsford, 2016), data structures developed to quickly query many files of data of short-read sequences from RNA-Seq data (and other data) for a particular sequence, were employed. These structures build on the concept of Bloom filters. The authors published software, which was subsequently Dockerized and wrapped in the Common Workflow Language (CWL) for use on the Seven Bridges Cancer Genomics Cloud pilot (Lau et al.; 2017). The supplemental file contains technical details about the methodology used.

### Ovarian Tumor Specimen Collection

Ovarian cancer samples were collected following procedures approved by the IRB from the Fred Hutchinson Cancer Research Center (FHCRC). Samples were (1) collected at initial debulking surgery using standardized protocols and (2) reviewed by a gynecological research pathologist to confirm the histological characteristics of the tissue; all tumor samples used in this article contained at least 70% malignant epithelium. Clinical data for RT-PCR screened samples are shown in Supplemental Table 4.

### RT-PCR Validation of fusions

Reverse transcription of RNA was performed (600 ng of each Ovarian cancer sample and 1 ug for neg. control HeLa and K562 total RNA) using Moloney Murine Leukemia Virus Reverse Transcriptase (M-MLV RT) enzyme (Promega) according to manufacturer’s recommendations. See Supplemental Table 4te for sample information. The reverse transcription was primed with equal parts of random N6 (PAN facility, Stanford University) at 2 .5 mM final concentration. cDNA reaction was diluted 1:10 and used 1 mL/10 mL PCR reaction and run for 40 cycles. Reactions were run on a 1x TBE 1.75% Agarose gel and imaged using Alpha Innotech AlphaImager™ (San Leandro, CA) gel imaging system. PCR-validated fusion transcripts were further confirmed using Sanger sequencing. PCR primers used and validated PCR sequences can be found below.

### Primers used and Sanger sequences obtained

For details and primers used, see Supplemental File.

## Acknowledgments

All RNA-Seq data was generated by The Cancer Genome Atlas project funded by the NCI and NHGRI. Information about TCGA and the investigators and institutions that constitute the TCGA Research Network can be found at https://cancergenome.nih.gov. We thank Rajat Rohatgi and Peter Wang for useful discussions and suggestions on the sMACHETE algorithm; Nathan Watson for kind help with implementing the MACHETE on the CGC; Peter Wang additionally for assistance with Figures; and Robert Bierman, Brandi Davis-Dusenbery and Kirsten Green for feedback on the manuscript. This work was supported by NCI grant R00 CA168987-03, NIGMS grant R01 GM116847, a JIMB seed grant, an NSF CAREER Award, McCormick-Gabilan Fellowship, and a Baxter Family Fellowship to J.S.. J.S. is an Alfred P. Sloan fellow in Computational & Evolutionary Molecular Biology. The Seven Bridges NCI Cancer Genomics Cloud pilot and work by EL were supported in part by the funds from the National Cancer Institute, National Institutes of Health, Department of Health and Human Services, under Contract No. HHSN261201400008C.

### Competing Interests

Erik Lehnert is an employee of Seven Bridges Genomics.

## List of Supplemental Tables

Supplemental Table 1: sMACHETE outputs from all analyzed Body Map and TCGA samples.

Note that, per correspondence with TCGA, we do not publish positions of the fusions found; these can be shared with researchers upon establishing, in conversation with TCGA, the correct protocols. Pan_cancer denotes the total number of samples in which the splice variant of the fusion is found by the SBT, summing over all SBTs. AbsPos1Pos2Diff is the absolute value of the difference between position 1 and position 2. MaxFreq denotes the frequency at which the splice variant appears in the SBT for the disease type. MaxCount is the number of samples in which the splice variant is found in the SBT for the disease. maxMAFreq and maxCompFreq refer, respectively, to the frequency of the splice variants as estimated by the SBT per disease type, in the discovery and test sets, respectively. Note that some fusions could be discovered in disease A and then found only in the test set for disease B.

Supplemental Table 2: Sample IDs and Metadata for Samples Analyzed with MACHETE

Supplemental Table 3: MACHETE outputs used as input to sMACHETE statistical models and SBT. Note that, per correspondence with TCGA, we do not publish any sample IDs or positions of the fusions found; these can be shared with researchers upon establishing, in conversation with TCGA, the correct protocols. AbsPos1Pos2Diff is the absolute value of the difference between position 1 and position 2.

Supplemental Table 4: List of ovarian tumor samples used for RT-PCR validation

Supplemental Table 5: Sample IDs and Metadata for Samples Analyzed with SBTs

### Supplemental Figures

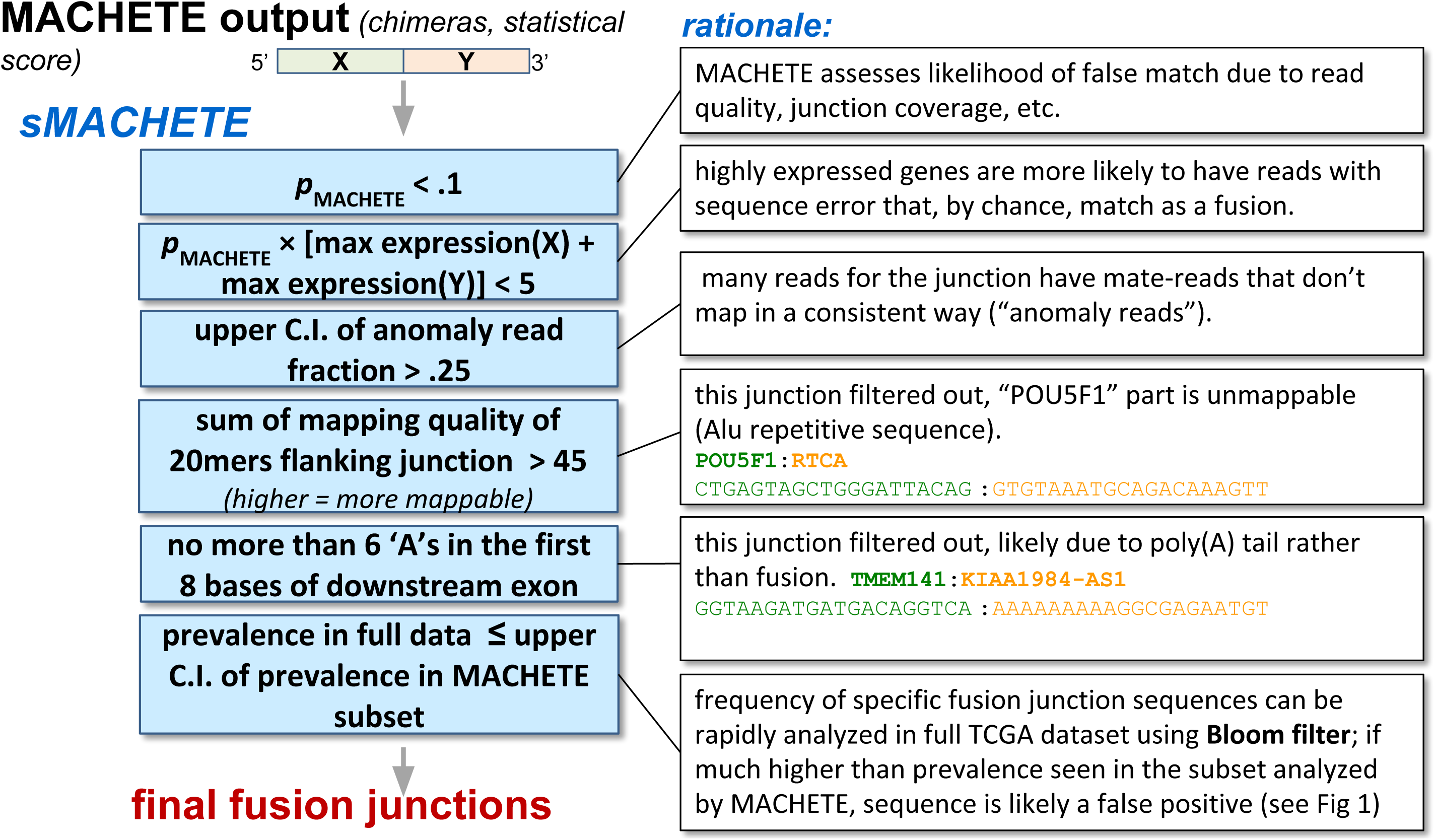

